# Neuraminidase-Based Cross-Protective Immunity against H5N1 Influenza Viruses in Humans

**DOI:** 10.64898/2026.06.07.730769

**Authors:** Iván Sanz-Muñoz, Carlos J. Ciria-Gil, Alejandro Martín-Toribio, Marina Toquero-Asensio, Javier Sánchez-Martínez, Carla Rodríguez-Crespo, Marta Hernandez, Iris Barragan-Martin, Sara Landeras-Bueno, Jose M. Eiros, Ahmed M. Elsayed, Luis Martinez-Sobrido, Aitor Nogales

## Abstract

Highly pathogenic avian influenza A(H5N1) virus continues to expand globally across wild birds, poultry, and multiple mammalian hosts, and the detection of genetic features associated with mammalian adaptation underscores the need to better define preexisting human immunity relevant to pandemic preparedness. Although serologic and vaccine studies have historically emphasized hemagglutinin (HA), neuraminidase (NA) is a key antigen capable of providing protective immunity. Here, we performed a retrospective analysis of NA-specific antibodies in human sera collected across ten influenza seasons spanning the pre- and post-2009 pandemic periods. A total of 749 paired human serum samples (1,498 total) were obtained prior to seasonal inactivated influenza vaccination (T1) and 28 days after vaccination (T2). NA-inhibiting antibodies were measured by enzyme-linked lectin assay (ELLA) using N1 antigens representing three genotypes (B.3.13, D1.1, and EA-2021-AB) from clade 2.3.4.4b A(H5N1) viruses. In parallel, HA-directed responses were assessed by hemagglutination inhibition (HAI). In addition, luciferase-based microneutralization assays were performed to assess the presence of neutralizing antibodies. NA-inhibiting antibodies were detectable against all three genotypes across seasons. Seasonal vaccination was associated with a modest but reproducible increase in NA-inhibiting titers, most apparent from the 2009-2010 season onward, coincident with the introduction of A(H1N1)pdm09 into seasonal vaccine formulations. In contrast, HAI activity against H5 was generally low or undetectable, despite detectable HA-reactive antibodies by ELISA. These data indicate that cross-reactive NA-specific antibodies are present in human sera and can be boosted by seasonal vaccination, supporting increased consideration of NA in influenza serosurveillance and vaccine evaluation.

**IMPORTANCE:** The continued spread of A(H5N1) viruses and reports of mammalian adaptation markers heighten concerns about potential zoonotic transmission into humans. Most studies of human antibody responses focus on influenza HA, but NA is also an important immune target. By analyzing paired sera from humans collected over ten influenza seasons, we show that NA-inhibiting antibodies cross-reacting with multiple contemporary genotypes from clade 2.3.4.4b A(H5N1) viruses are detectable and can be modestly boosted seasonal inactivated influenza vaccination, particularly after the 2009 pandemic. These findings highlight the importance of inducing NA-specific antibodies during influenza vaccination and in including NA in surveillance and in the assessment of vaccines intended to improve preparedness for emerging influenza viruses.

## INTRODUCTION

Influenza A viruses (IAVs) are enveloped, negative-sense, single-stranded RNA viruses in the *Orthomyxoviridae* family with zoonotic potential to both human and animal health (1, 2). IAVs primarily circulate in avian reservoirs, yet their exceptional evolutionary plasticity facilitates the emergence of variants capable of infecting mammals and, in some cases, spreading efficiently among humans with epidemic or pandemic consequences (1–6).

Two major viral surface glycoproteins, hemagglutinin (HA) and neuraminidase (NA) are embedded in the viral lipid envelope and form the basis for subtype classification into 18 HA (H1-H18) and 11 NA (N1-N11) IAV subtypes (1–6). Importantly, HA, and to a lesser extent NA, are the principal targets of neutralizing antibodies (NAbs) induced by vaccination and/or by natural infection (6–8). Humans are repeatedly exposed to seasonal IAV subtypes H1N1 and H3N2, as well as influenza B virus (IBV), through recurrent infections and/or routine vaccinations (5). Moreover, individuals born before 1968 are likely to have been exposed to both H1N1 and H2N2 IAV subtypes, while those born after 1968 have been probably exposed to H1N1 and H3N2 (9).

Vaccination remains the most effective strategy to reduce morbidity and mortality associated with IAV infections in both humans and animals (2, 3, 10–13). Inactivated influenza vaccines (IIVs) primarily elicit strain-specific humoral responses directed against the viral HA (7, 8), and therefore, limited protection against antigenically distinct drifted seasonal or shifted pandemic strains (14). Notably, several studies have reported heterotypic antibody responses following seasonal vaccination, including antibodies capable of recognizing diverse avian influenza viruses (AIVs) (15, 16). Nevertheless, the extent to which seasonal IIV-induced immunity confers meaningful protection against heterologous IAV subtypes in humans remains unclear (17–19).

Highly pathogenic avian influenza (HPAI) H5N1 viruses are no longer a distant zoonotic threat but an increasingly pervasive feature of the global infectious disease landscape (4, 5). Their unprecedented expansion across wild avian reservoirs, poultry systems, and a widening range of mammalian hosts signals a profound ecological shift with direct implications for human health (4, 6). The recent emergence of H5N1 genotype 2.3.4.4b, including variants carrying molecular signatures of mammalian adaptation, reinforce concerns that the next influenza pandemic may arise from viruses already in active global circulation (20, 21). In this setting, defining the extent and nature of pre-existing human immunity is not only a scientific priority but a prerequisite for effective pandemic preparedness (22).

For decades, the field has been dominated by a HA-centric paradigm, in which NAbs targeting viral entry are regarded as the principal mediators of protection (3, 23). This framework has shaped both serological surveillance and vaccine design, with hemagglutination inhibition (HAI) titers serving as the main canonical correlate of immune protection for human vaccines. However, NA, the viral enzyme responsible for particle release from infected cells, represents a critical but comparatively neglected antigenic target (24–27). A growing body of evidence indicates that NA-specific antibodies can reduce viral replication, limit transmission, and confer protection even in the absence of strong HA-directed neutralization (28–30). Even more, NA is more conserved than HA, thus broad immune responses are probably more frequent. Despite these insights, NA immunity remains largely unquantified and systematically overlooked (31). In addition, data on baseline human NA cross-reactivity to contemporary H5N1 are limited, and it remains to be determined whether seasonal influenza vaccines induce anti-NA responses to avian NAs over time, especially across the antigenic shift surrounding the 2009 pandemic.

This blind spot is particularly consequential in the context of antigenically novel viruses such as H5N1 (**3, 32**). Human populations are typically immunologically naïve at the level of HA but may harbor layers of cross-reactive immunity that are not captured by conventional assays (3, 25, 27, 32). Whether NA-directed responses constitute a meaningful component of this hidden immunity, and whether they can be shaped by prior infections or vaccination, remains an open and largely unexplored question (26, 31). Addressing this gap is essential for establishing correlates of protection and toward a more comprehensive understanding of influenza immunity (23).

Here, we interrogate a decade of human serological data to define the prevalence, magnitude, and inducibility of NA-specific antibodies against representative genotypes of circulating H5N1 clade 2.3.4.4b strains. By integrating longitudinal sampling before and after seasonal influenza vaccination with complementary assays of NA and HA immunity, we uncover evidence of widespread yet underappreciated NA-directed responses in human populations. Collectively, these results emphasize the importance of integrating NA into both immunological surveillance frameworks and vaccine design efforts, with direct relevance to pandemic preparedness in the context of increasing zoonotic spillover.

## MATERIAL AND METHODS

### Study design and sample set

We designed a prospective study including adults and elderly people who were vaccinated with the IIV during the 2006-2007, 2007-2008, 2008-2009, 2009-2010, 2010-2011, 2011-2012, 2012-2013, 2015-2016, 2016-2017, and 2017-2018 influenza seasons. A total of 749 paired sera (total of 1,498 samples) were evaluated. We recruited individuals from three different settings. The first group included workers at two automobile factories of the Renault Group in Spain in the cities of Valladolid and Palencia. The second group included individuals in Spain living in two nursing homes in the province of Valladolid, under the supervision of the “Diputación de Valladolid” Council. The third one included individuals vaccinated in primary care in different settings in the community of Castilla y León (Spain). A serum sample from each individual was obtained at two different time points. The first one was just before being vaccinated with the seasonal IIV (T1). After that, a new serum sample was obtained 1 month after vaccination (T2). With this study design, we aimed to determine the immunological background of antibodies against H5N1, and the heterotypical response elicited by seasonal vaccines (**Figure 1) (Table1**). During these seasons, multiple trivalent/quadrivalent standard-dose, adjuvanted and high dose IIV were used (**Table 1**). The IIV administered contained the IAV and IBV strains recommended by the World Health Organization (WHO) for each influenza season for the Northern hemisphere. Selected samples were positive by HAI for IAV pH1N1 and H3N2 at post-vaccination stage at 83.8% and 76.4% respectively, and seroconversion was observed in 13.5% for pH1N1 and 33.5% for H3N2 of individuals. This work was approved by the Ethics Committee of the Eastern Health Area of Valladolid under three different codes (Cod: PI 21-2314, PI 21-2442 and PI 22-2894). All participants provided written informed consent prior to vaccination and sampling. Additionally, this research was conducted according to the Declaration of Helsinki.

**Figure 1.**
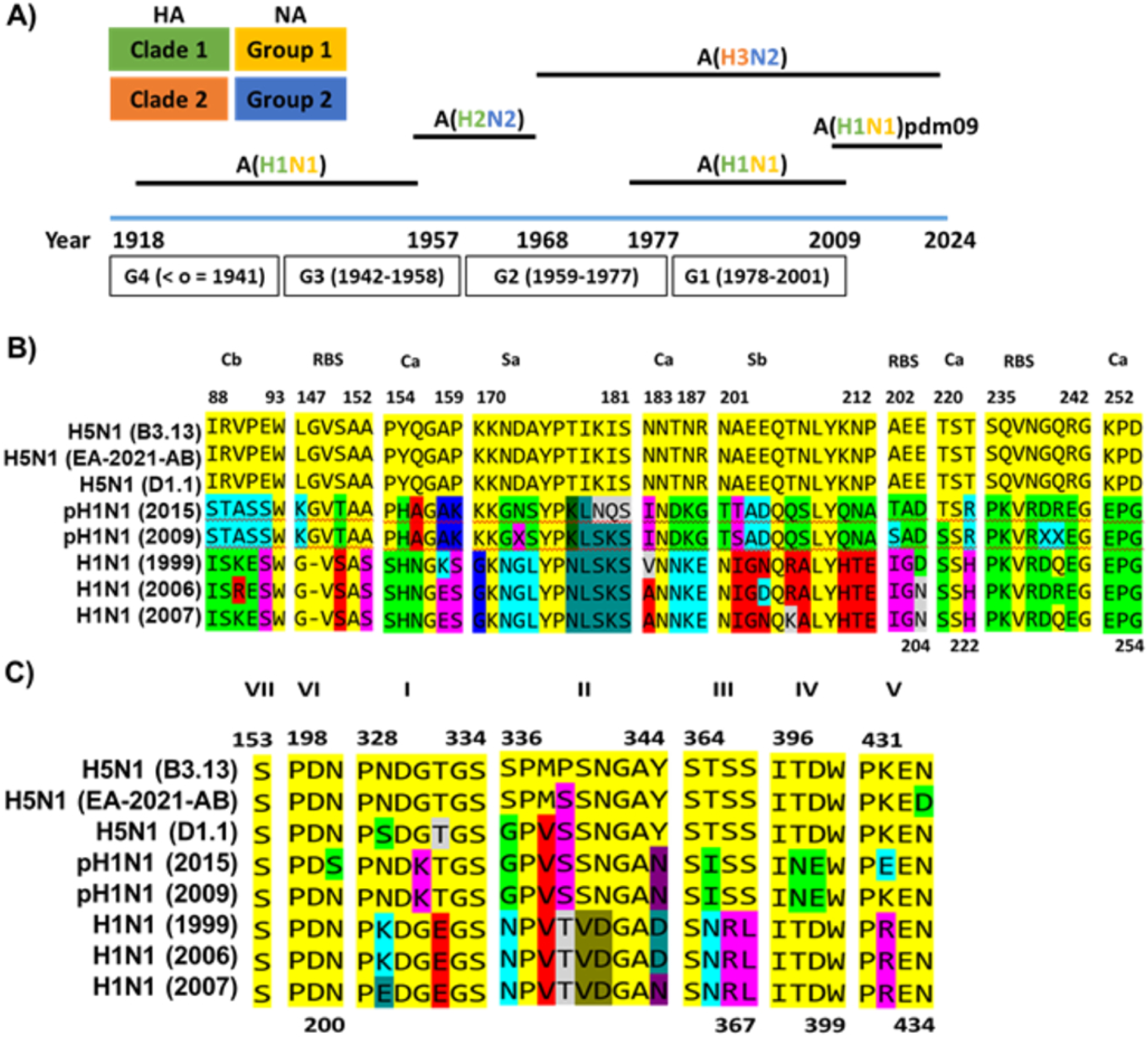
Historical circulation and antigenic-site divergence between human H1 and avian H5 IAV. (**A**) Temporal dynamics of the circulation and co-circulation (1918–2024) of distinct IAV subtypes in humans, with HA clades and NA groups indicated. (**B**) Multiple-sequence alignment of the HA head antigenic regions showing amino acid differences between H5 and H1 strains in the canonical antigenic (Ca, Cb, Sa, Sb) and within the receptor-binding (RBS) sites. (**C**) Multiple-sequence alignment of putative NA antigenic sites. Seasonal H1N1 and avian H5N1 strains show the greatest divergence, whereas the 2009 pH1N1 displays an intermediate pattern. H5N1: B3.13 A/Texas/37/2024; EA-2021AB A/chicken/Egypt/F71-F114C/2022; D1.1 A/ Louisiana/12/2024. pH1N1: A/Michigan/45/2015; A/California/7/2009. H1N1: A/New Caledonia/20/99; A/Solomon Islands/3/2006; A/Brisbane/59/2007.

**Table 1.**
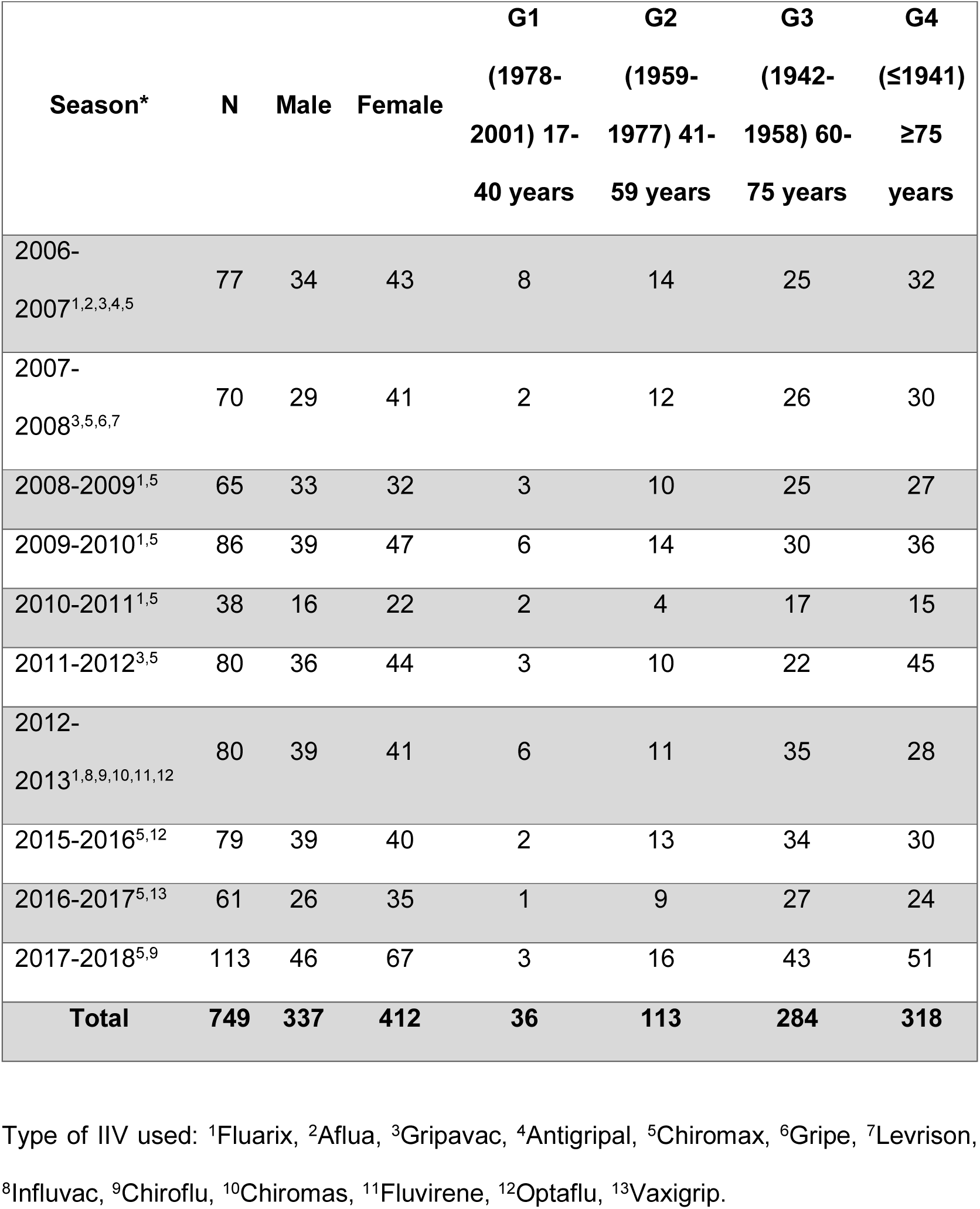
Cohort description and epidemiological characteristics.

### Cell lines

Human embryonic kidney 293T (HEK293T; ATCC CRL-11268) and Madin-Darby canine kidney (MDCK; ATCC CCL-34) cells were grown in Dulbecco’s modified Eagle’s medium (DMEM) supplemented with 5% fetal bovine serum (FBS) and 1% penicillin (100 units/mL)-streptomycin (100 µg/mL)-2 mM L-glutamine (PSG) at 37°C in air enriched with 5% CO_2_.

### Rescue of recombinant viruses and viral infections

Recombinant IAVs were rescued using previously described ambisense reverse genetics approaches (11, 16, 33, 34). Briefly, co-cultures (1:1) of HEK293T/MDCK cells (twelve-well plate format, duplicates) were co-transfected, using LPF3000 (Invitrogen), in suspension with six PR8 ambisense pHW-PB2, -PB1, -PA, -NP, -M, and -NS wild-type (WT) or Nanoluciferase (Nluc) plasmids and the pHW-HA and -NA from A/Texas/37/2024 (H5N1, HPAIV clade 2.3.4.4b, genotype B3.13) and A/ Louisiana/12/2024 (H5N1, HPAIV clade 2.3.4.4b, genotype D1.1). In both cases, the H5 polybasic cleavage site motif was changed to a monobasic cleavage site. The recovered viruses were plaque purified and propagated on MDCK cells at 37°C. After viral infection, cells were maintained in DMEM supplemented with 0.3% bovine serum albumin (BSA), 1% PSG, and 1 µg/mL tosyl-sulfonyl phenylalanyl chloromethyl ketone (TPCK)-treated trypsin (Sigma) (16, 33, 35, 36). Recombinant (r)WT and Nluc-expressing viruses encoding the HA and NA of A/chicken/Egypt/F71-F114C/2022 (H5N1, HPAIV clade 2.3.4.4b, genotype EA-2021-AB) or A/Anhui/1/2013 (H7N9, LPAIV) were previously described (16).

### Virus inactivation using beta-propiolactone (BPL)

BPL was combined with 10 mL of virus suspension (1×10^7^ plaque-forming unit (PFU)) and 0.5 mL of NaHCO_3_ (7.5%) to attain a final concentration of 0.05%. This buffered BPL/virus combination was then thoroughly vortexed to form a homogenous mixture and incubated at 4°C for 16 h in an orbital shaker. The preparation was subsequently incubated at 37°C for 2 h for hydrolysis of BPL (16, 37, 38). After completion of treatment, the virus aliquots were stored at -80°C until further use and inactivation was confirmed via three blind-passages and plaque assay (16).

### Enzyme-linked immunosorbent assay (ELISA)

HA-, NA-, and NP-specific serum IgG antibodies were detected by ELISA (16). Briefly, 96-well plates were coated for 16 h at 4°C with 100 ng per well of recombinant H5 (A/bald Eagle/Florida/125/2017 H5N1; NR-59424, Bei Resources), N1 (A/Texas/37/2024 H5N1; 41016-V08B, Sino Biologicals), and NP (A/Wisconsin/67/2022 H1N1 or A/Nanjing/1/2015 H5N1) proteins. After washing with PBS, the coated wells were blocked with PBS-T (Tween, 0.05%) containing 5% BSA and then incubated with a two-fold dilution (starting dilution 1:100) of human serum at 37°C. After 1 h of incubation, the plates were washed with PBS-T and incubated with HRP-conjugated goat anti-human IgG for 1 h at 37°C. The reactions were developed with tetramethylbenzidine substrate (BioLegend) for 10 min at room temperature, quenched with 2 N H_2_SO_4_, and read at 450 nm. Serum samples from uninfected mice or mice infected with Rift Valley Fever Virus from a previous experiment were used as internal negative controls (39). We used as threshold for positive samples the average +3 SD of the negative control.

### Hemagglutination inhibition (HAI) assays

HAI assays were performed at the National Influenza Center of Valladolid (NICV) in BSL2 facilities. HAI assays were used as reference to analyze antibody responses induced by influenza vaccines (40). To that end, viruses were first inactivated with BPL. Serum samples were centrifuged at 2,500 rpm for 10 min, and the supernatants were stored at -20°C. For each serum sample, 100 µL were added to 300 µL of receptor-destroying enzyme (RDE; Denka Seiken, Tokyo, Japan) to remove nonspecific HA inhibitors. Following the manufacturer’s instructions, this mixture was incubated at 37°C in a water bath for 12–18 h, followed by enzyme inactivation for 1 h at 56°C. To perform HAI, 50 µL serial two-fold dilutions of each serum (starting dilution of 1:10) were made in 96 microwell V plates, and then 50 µL of standard viral samples containing four units of HA were added to each well and incubated for 30 min at room temperature. Next, 50 µL of 0.5% chicken erythrocytes were added and plates were incubated at room temperature for another 30 min. The titer of antibodies was defined as the highest dilution that completely inhibited hemagglutination. The limit of detection of the assay was 1:10. When the antisera titer was below a detectable threshold, due to a shortage or lack of antibodies, it was conventionally expressed as 5, which is half of the lowest detection threshold.

### Enzyme-linked lectin assays (ELLA)

NA activity and inhibition were measured using ELLA adapted from previously described methods (41–43). High-binding 96-well plates (MaxiSorp, Thermo Fisher Scientific) were coated with fetuin (25 µg/ml, 100 µl per well; Sigma-Aldrich) in PBS and incubated at 4°C overnight. Plates were washed three times with PBS containing 0.05% Tween-20 (PBS-T) prior to use. For determination of NA activity, virus was serially diluted in PBS supplemented with CaCl₂ (20 mM), MgCl₂ (20 mM), 1% PBS) and 0.05% Tween-20 and added to fetuin-coated plates. Following incubation at 37°C for 16-18 h, plates were washed five times with PBS-T. Peanut agglutinin conjugated to horseradish peroxidase (PNA–HRP; Sigma-Aldrich) diluted 1:1,000 in PBS containing CaCl₂, MgCl₂ and 1% BSA was added (50 µl per well) and incubated for 2 h at room temperature in the dark. Plates were washed 5x with PBS-T and developed with TMB substrate (Thermo Fisher Scientific) for 10-15 min before stopping the reaction with 1N H₂SO₄. Absorbance was measured at 450 nm. The effective concentration of virus corresponding to 90–98% of maximal signal (EC₉₀ or EC₉₈) was determined by nonlinear regression analysis (GraphPad Prism). For inhibition assays, sera were serially diluted in PBS and mixed 1:1 with virus at 2xEC₉₀/EC₉₈ directly in fetuin-coated plates. Plates were incubated at 37°C for 16–20 h, and NA activity was detected as described above. Background-subtracted absorbance values were normalized to virus-only controls to calculate percent NA activity (% NA), and percent inhibition (% NI) was defined as 100 - % NA. Each assay included background controls, virus-only controls, and positive and negative reference sera.

### Reporter-based microneutralization (MN) assays

Nluc-based MN assays were performed as previously described (16). Briefly, human serum samples were two-fold serially diluted, starting at a 1:200 dilution. Subsequently, 200 PFU of NLuc-expressing H5N1 virus were added to each serum dilution, and the mixtures were incubated for 1 h at room temperature. MDCK cells seeded in 96-well plates (5 x 10⁴ cells per well, triplicates?) were then infected with the serum-virus mixtures and incubated for 48 h at 37°C. NLuc activity in culture supernatants was quantified using the Nano-Glo luciferase substrate (Promega) and measured with a Lumicount luminometer. In an alternative assay format, MDCK cells were first infected with 200 PFU of NLuc-expressing H5N1 viruses. After 1 h of viral adsorption, the inoculum was removed and replaced with fresh medium containing serial two-fold dilutions of human sera (starting at 1:200). Cells were subsequently incubated for 48 h at 37°C, after which NLuc activity in the culture supernatants was measured as described above. NLuc values of virus-infected cells in the absence of antibody were used to calculate 100% viral infection. Cells in the absence of viral infection were used to calculate the luminescence background. After the NLuc activity was measured, cells were fixed and stained with crystal violet solution.

### Statistical analysis

The statistical analysis was focused on a descriptive and inferential study of the antibody titers based on the age of the patients. The results were analyzed using the classical serological parameters of the European Medicines Agency for the evaluation of vaccine efficacy (44, 45). These parameters included the seroprotection rate (SPR), seroconversion rate (SCR), and geometric mean titer (GMT). Seroprotection was considered when an HAI titer of 1/40 was achieved (46), and seroconversion was defined as a titer increase of at least fourfold between pre- and post-vaccination sera. In addition, seroconversion was considered to have occurred in cases of negative pre-vaccination sera that achieved 1/40 titers after vaccination. The use of 1/40 titer as a seroprotection titer was based on the previous knowledge and consolidated titers against seasonal influenza viruses (46, 47). It was also based on previous publications suggesting that this 1/40 titer can be used for both seasonal and avian viruses for HAI analysis as well as MN assays (48, 49). Moreover, phase I and II clinical studies conducted to evaluate immunogenicity of some avian influenza vaccines for humans use the 1/40 titer as a surrogate of protection for HAI, based on the CBER/CHMP (Center for Biologics Evaluation and Research/Committee for Medical Products for Human Use) criteria (48, 50). For the analysis, a negative value of HAI titers was presented as “5” (taking 1/10 as the lower detection threshold) to avoid errors during the logarithm data transformation for calculation of GMTs. The values are presented as the means (95% CIs). The volunteer characteristics, SPR and SCR were compared using χ2 tests. GMTs were compared using Student’s t-test. The data from the neutralization assays were compared using two-way analysis of variance (ANOVA). Data analysis was performed using GraphPad Prism and Microsoft Excel. A P-value <0.05 was considered to indicate statistical significance.

## RESULTS

### Cohort description and sample distribution

A total of 749 participants were recruited for this study and paired pre-vaccination (T1) and post-vaccination (T2) serum samples (1,498 sera in total) were collected (**Table 1**). Specimens were collected across 10 influenza seasons spanning 2006–2007 to 2017–2018 (with no collections shown for 2013-2014 and 2014-2015), with seasonal sample sizes ranging from 38 to 113 (median 78, mean 74.9 participants/season). Overall, 337/749 (45.0%) participants were male and 412/749 (55.0%) were female. Participants were grouped by birth cohort/age at vaccination as follows: G1 (born 1978-2001; 17-40 years), G2 (1959-1977; 41-59 years), G3 (1942-1958; 60-75 years), and G4 (≤1941; ≥75 years). The study population was predominantly older adults, with G4 comprising 318/749 (42.5%), G3 284/749 (37.9%), G2 113/749 (15.1%), and G1 36/749 (4.8%) of participants, indicating that ∼80% of paired samples came from individuals aged ≥60 years. We selected four different age groups for this study related with the pandemics and influenza emergences of the XX-XXI century (**Fig 1A**).

### Determining human immunity against H5N1 before and after IIV

HAI titers were measured for each subject against three clade 2.3.4.4b H5N1 genotypes (B.3.13, D1.1, and EA-2021-AB) using inactivated virus. HAI titers were low or undetectable in sera collected before vaccination (T1) and after receipt of the IIV (T2), with only a small number of individuals reaching protective titers (≥1:40) against at least one H5N1 genotype (data not shown), consistent with our recent report (16) and previous studies (16, 51–56). Most of the subjects showing protective titers were found against D.1.1. H5N1 virus (9 out of 749; 1.2%). In contrast, seroconversion and seroprotection rates were moderate to high against pH1N1 and H3N2 viruses in all selected samples as previously described (57–59). Substantial amino-acid differences between H5N1 and H1N1 or pH1N1 HA, particularly across the conventional head antigenic sites (Ca, Cb, Sa, Sb) and the receptor-binding site (RBS) (**Figure 1B**), may account for these results and contribute to the reduced HAI reactivity (low titers) against H5N1, even in post-vaccination samples.

In addition, the levels at the population level of IgG antibodies recognizing H5, N1_H5N1_ or NP_H5N1_ and NP_pH1N1_ viral antigens were measured by ELISA using purified recombinant proteins (**Figure 2**) across the ten influenza seasons. Population IgG binding to H5 (**Figure 2A**), N1 (**Figure 2B**), and NP (**Figure 2C**) was observed both before (T1) and after vaccination (T2). For H5 and N1, most seasons showed a modest increase in IgG levels at T2, suggesting limited induction of cross-reactive anti-HA and/or anti-NA antibodies following administration of the seasonal IIV. Notably, the observed anti-H5 and anti-N1 reactivity may reflect recognition of conserved epitopes within HA and NA. In contrast, IgG levels against NP_H5N1_ (**Figure 2C top**) or NP_pH1N1_ (**Figure 2C bottom**) were comparable at T1 and T2, indicating that NP-specific antibodies were not boosted by seasonal IIV. However, these responses could be important for cellular mediated responses. Given that, a large proportion of the population has experienced prior IAV exposure through infection, vaccination or both (often repeatedly), the presence of baseline antibodies to these antigens was expected.

**Figure 2.**
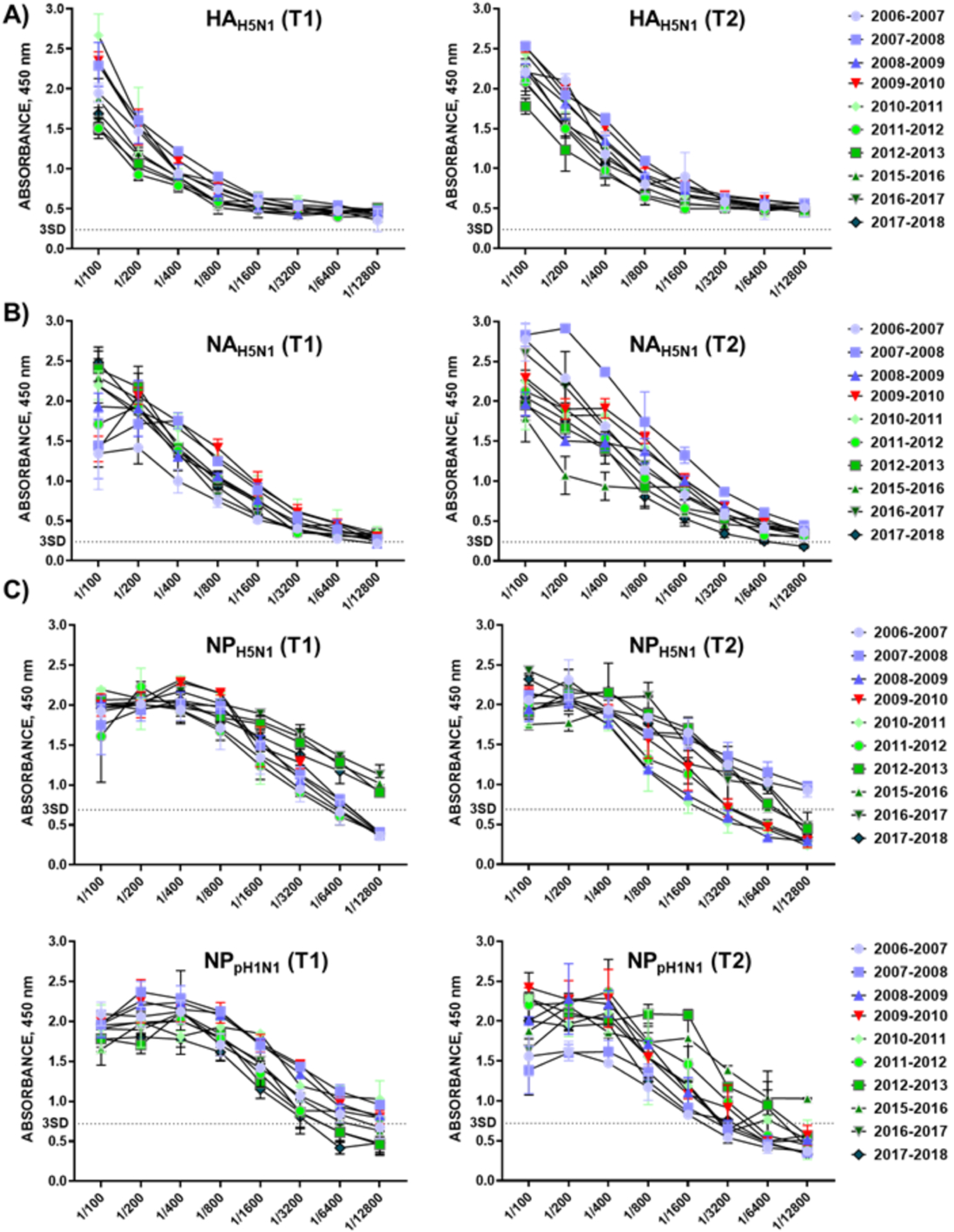
IgG binding to H5N1 HA, NA, and NP in human collected across ten influenza seasons. Serum IgG reactivity to recombinant HA (**A**) and NA (**B**) of H5N1, and NP of H5N1 or pH1N1 (**C**) was measured by ELISA across ten influenza seasons using purified recombinant proteins. HA_H5N1_: A/bald Eagle/Florida/125/2017 H5N1 NA_H5N1_: A/Texas/37/2024 H5N1; NP_H5N1_: A/Nanjing/1/2015; NP_pH1N1_: A/Wisconsin/67/2022. Antibody binding was assessed at baseline (T1) and 28 days post-vaccination (T2). Curves represent absorbance at 450 nm across serial serum dilutions; the dotted line indicates the 3 SD threshold.

### Baseline cross-reactive NA antibodies to H5N1 genotypes

Because HA-directed HAI activity against H5N1 was low, we quantified NA-inhibiting antibodies, as NA-inhibiting immunity can limit viral spread and disease and may be detectable even when HA is antigenically mismatched. Since H5N1 and human H1N1 or pH1N1 viruses share N1 (group 1) NA, cross-inhibitory anti-N1 responses in human sera could be anticipated. Therefore, we assessed the presence of NA-inhibiting antibodies in the 749 paired human serum samples; each analyzed individually using the ELLA. NA-inhibiting activity was evaluated across the 10 influenza seasons included in the study using sera collected before (T1) and after vaccination (T2). We focused on H5N1 clade 2.3.4.4b genotype B3.13 because of its similarity at the level of NA antigenic sites (**Figure 1C**) and overall NA protein sequence (**Table 2**). When visualized as violin plots, the data showed consistently high NA-inhibiting antibody levels across all seasons (**Figure 3A**). Notably, vaccination increased NA-inhibiting activity, with the largest post-vaccination rise observed in the season following the pH1N1 event (**Figure 3A**). This enhanced response may reflect the greater similarity between the N1 of H5N1 and the N1 of pH1N1 compared with the N1 of seasonal H1N1 viruses circulating prior to the pH1N1 responsible of the 2009 pandemic (**Fig 1C** and **Table 2**). In addition, IIV increased NA-inhibiting antibody levels across all age groups (**Figure 4A**). However, older age groups exhibited higher NA-inhibiting antibody levels overall (**Figure 4B**), potentially reflecting cumulative exposure to a broader diversity of NA variants through repeated infection and/or vaccination. Although we assessed whether anti-NA antibody levels differed by sex, we observed no significant differences between males and females at either T1 or T2. In addition, we also assessed NA-inhibiting antibodies against all H5N1 clade 2.3.4.4b genotypes at the population level using season-specific pooled sera (**Figure 5**). ELLA revealed similarly high NA-inhibiting activity across the three clade 2.3.4.4b H5N1 genotypes tested (B3.13, D1.1, and EA-2021-AB), reflecting their sequence similarity (**Figure 1C**).

**Figure 3.**
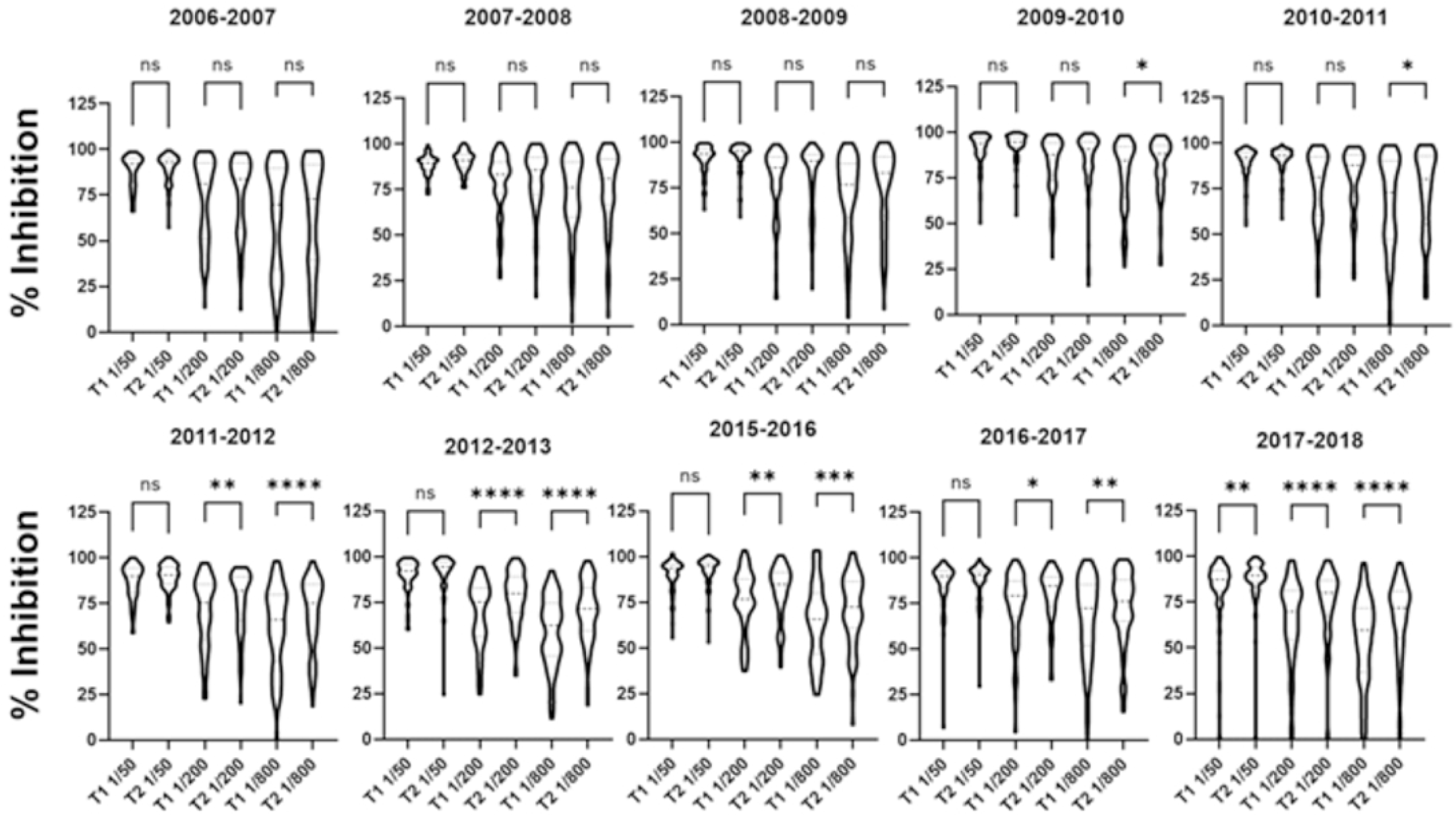
NA-inhibiting antibody responses to H5N1 (genotype B3.13) measured by ELLA across ten influenza seasons. NA inhibition is shown as % inhibition for each season, with individual values displayed as violin plots (T1: baseline; T2: post-vaccination). Statistical significance is indicated above brackets (two-way ANOVA): ns, not significant; P < 0.05; P < 0.01; P < 0.001; P < 0.0001.

**Table 2.** Comparative NA Sequence Identity Among H5N1, H1N1, and pH1N1.

|  | **A/NC/99**<br>**H1N1** | **A/SI/06**<br>**H1N1** | **A/Bris/07**<br>**H1N1** | **A/Cal/2009**<br>**pH1N1** | **A/Mic/2015**<br>**pH1N1** | **EA-2021AB H**<br>**5N1** | **B3.13**<br>**H5N1** | **D1.1**<br>**H5N1** |
| --- | --- | --- | --- | --- | --- | --- | --- | --- |
| **A/NC/99**<br>**H1N1** | **100** | 97.2 | 97.7 | 79.3 | 78.4 | 62.2 | 61.8 | 61.6 |
| **A/SI/06**<br>**H1N1** | 97.2 | **100** | 98.4 | 79.1 | 78.4 | 61.2 | 60.7 | 60.9 |
| **A/Bris/07**<br>**H1N1** | 97.7 | 98.4 | **100** | 78.8 | 77.9 | 61.7 | 61.2 | 61.4 |
| **A/Cal/2009**<br>**pH1N1** | 79.3 | 79.1 | 78.8 | **100** | 96.6 | 61.7 | 61.3 | 61.1 |
| **A/Mic/2015**<br>**pH1N1** | 78.4 | 78.4 | 77.9 | 96.6 | **100** | 61.7 | 61.3 | 61.1 |
| **EA-2021AB H**<br>**5N1** | 62.2 | 61.2 | 61.7 | 61.7 | 61.7 | **100** | 97.3 | 97.7 |
| **B3.13**<br>**H5N1** | 61.8 | 60.7 | 61.2 | 61.3 | 61.3 | 97.3 | **100** | 98.4 |
| **D1.1 H5N1** | 61.1 | 60.9 | 61.4 | 61.1 | 61.1 | 97.7 | 98.4 | **100** |

**Figure 4.**
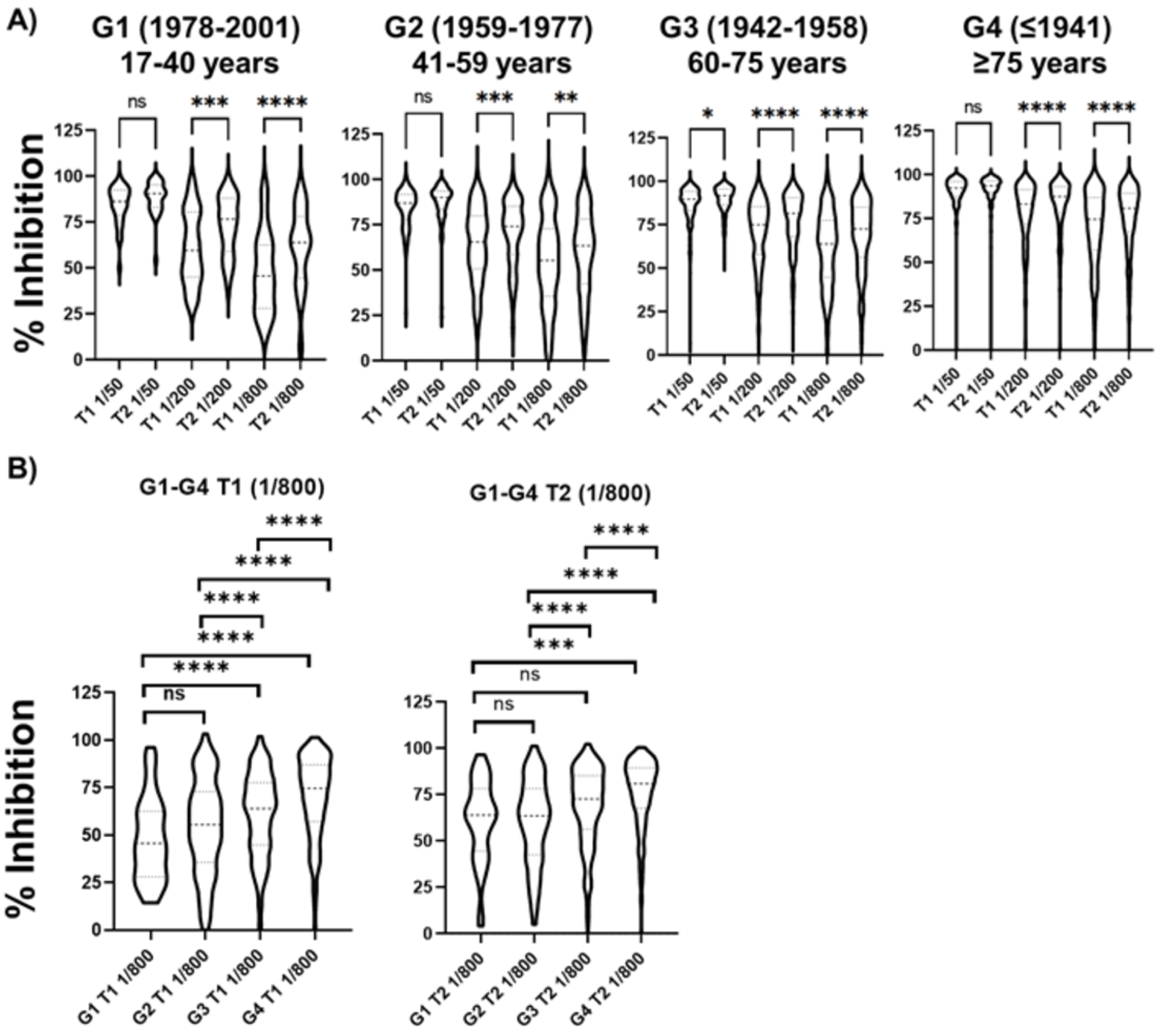
Age-associated NA-inhibiting antibody responses to H5N1 (genotype B3.13). (**A**) NA-inhibiting antibody activity (ELLA; % inhibition) was evaluated at baseline (T1) and 28 days post-vaccination (T2) across four age groups: G1 (1978–2001; 17–40 years), G2 (1959–1977; 41–59 years), G3 (1942–1958; 60–75 years), and G4 (≤1941; ≥75 years). (**B**) Group comparisons of NA inhibition at 1:800 serum dilution are shown for T1 (left) and T2 (right) across G1–G4 (**Table 1**). Significance is indicated above brackets (two-way ANOVA): ns, not significant; P < 0.05; P < 0.01; P < 0.001; P < 0.0001.

**Figure 5.**
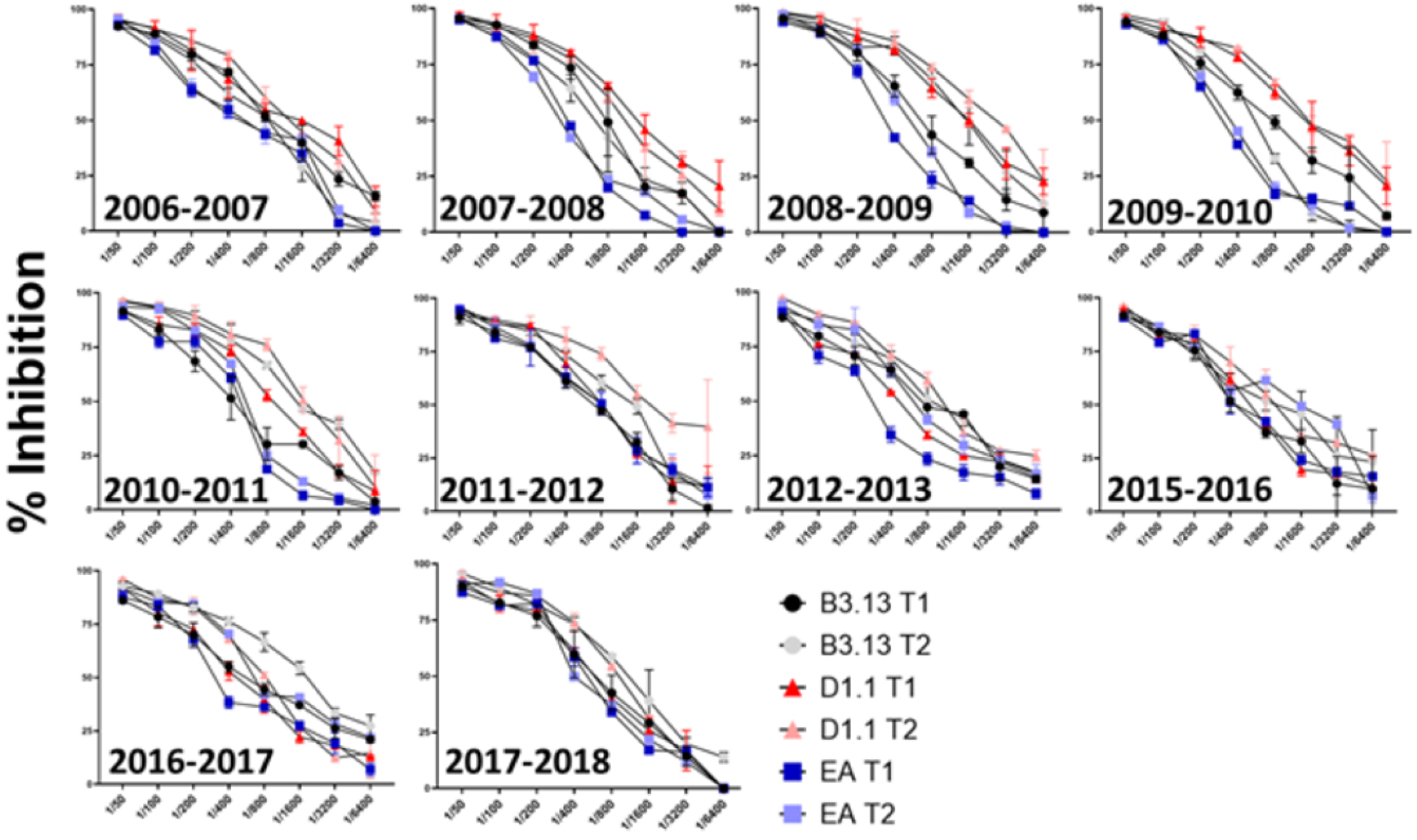
NA-inhibiting antibody responses to H5N1 across multiple influenza seasons. NA-inhibiting activity was quantified by ELLA using serial serum dilutions at baseline (T1) and post-vaccination (T2) for each influenza season (2006–2007 to 2017–2018, as indicated). For each season, inhibition curves are shown against NA from three H5N1 genotypes: B3.13 (black/gray circles), D1.1 (red/pink triangles), and EA-2021-AB (dark/light blue squares). The y-axis indicates % inhibition, and the x-axis indicates the reciprocal serum dilution. Points are displayed with error bars to indicate variability among samples.

### Cross-reactive antibodies to neutralize IAV H5N1 *in vitro*

NAbs represent a key immunological correlate of protection following influenza vaccination or infection (60–64). However, conventional neutralization assays are often labor-intensive and typically rely on secondary readouts of viral replication (e.g., cytopathic effect or immunostaining), which limits throughput and scalability. We previously showed that this constraint could be mitigated by using IAV engineered to express fluorescent or luciferase reporter genes, enabling direct monitoring and quantification of influenza infection (16, 33, 60). Building on this approach, we generated PR8(H5N1)-NLuc reporter viruses encoding the HA and NA of the clade 2.3.4.4b H5N1 genotypes evaluated in this study (B3.13, D1.1, and EA-2021-AB), following our prior strategy (16). We then used these reporter viruses to address two related questions across the 10 influenza seasons included in the study: (i) the extent of pre-existing population immunity against H5N1 (T1), and (ii) whether seasonal IIV can elicit heterosubtypic responses capable of neutralizing H5N1 (T2). Serum samples collected before (T1) and after (T2) vaccination (**Table 1**) were tested for their ability to neutralize PR8(H5N1)-NLuc viruses using an NLuc-based MN assay (16, 33). Using the standard NLuc-based MN format, where virus and human sera are pre-incubated before infection of MDCK cells, we did not detect neutralization of any of the three H5N1 genotypes at either T1 or T2 in any of the sera collected at the 10 seasons (data not shown). This lack of detectable neutralization is consistent with our HAI results and with our previous observations (16), supporting the conclusion that antibodies elicited by seasonal IIV generally do not block H5 HA-mediated entry in HAI or classical MN assays.

Notably, when we modified the NLuc-based MN assay by adding human sera after viral infection (supplementing sera in the post-infection culture supernatants), we observed measurable neutralization across all seasons at both T1 and T2 (**Figure 6**). Importantly, neutralization levels were comparable across the three H5N1 reporter viruses, and the highest neutralization titers were observed for cohorts from the 2007-2012 seasons. Because this modified MN assay is expected to better capture inhibitory activities that act after viral entry, including antibodies that restrict viral egress and cell-to-cell spread (e.g., anti-NA antibodies) and, potentially, antibodies targeting conserved HA stalk epitopes that can limit subsequent rounds of infection, these findings suggest that heterosubtypic functional antibody activity is present but is not readily detected by assays focused on pre-entry neutralization. Taken together, results from the post-infection MN assays, in conjunction with the ELLA results, support that antibodies against NA N1 contribute, at least in part, to the neutralization levels observed with H5N1 viruses. This interpretation is consistent with the notion that NA-directed antibodies may not prevent initial infection but can substantially reduce the production and release of progeny virions (8, 24, 26). The elevated neutralization observed in 2007-2012 may reflect temporal differences in exposure histories and/or seasonal vaccine-driven boosting of cross-reactive NA immunity; a possibility warranting further investigation. Differences in NA functional activity and in the induction of anti-NA responses have been reported between different vaccine types (e.g., split versus subunit or adjuvanted formulations), indicating that vaccine brand and formulation may contribute to the heterogeneity in NA inhibition observed across seasons. Thus, inter-seasonal variation in NA inhibition should be interpreted in the context of both population immunological history and vaccine product characteristics.

**Figure 6.**
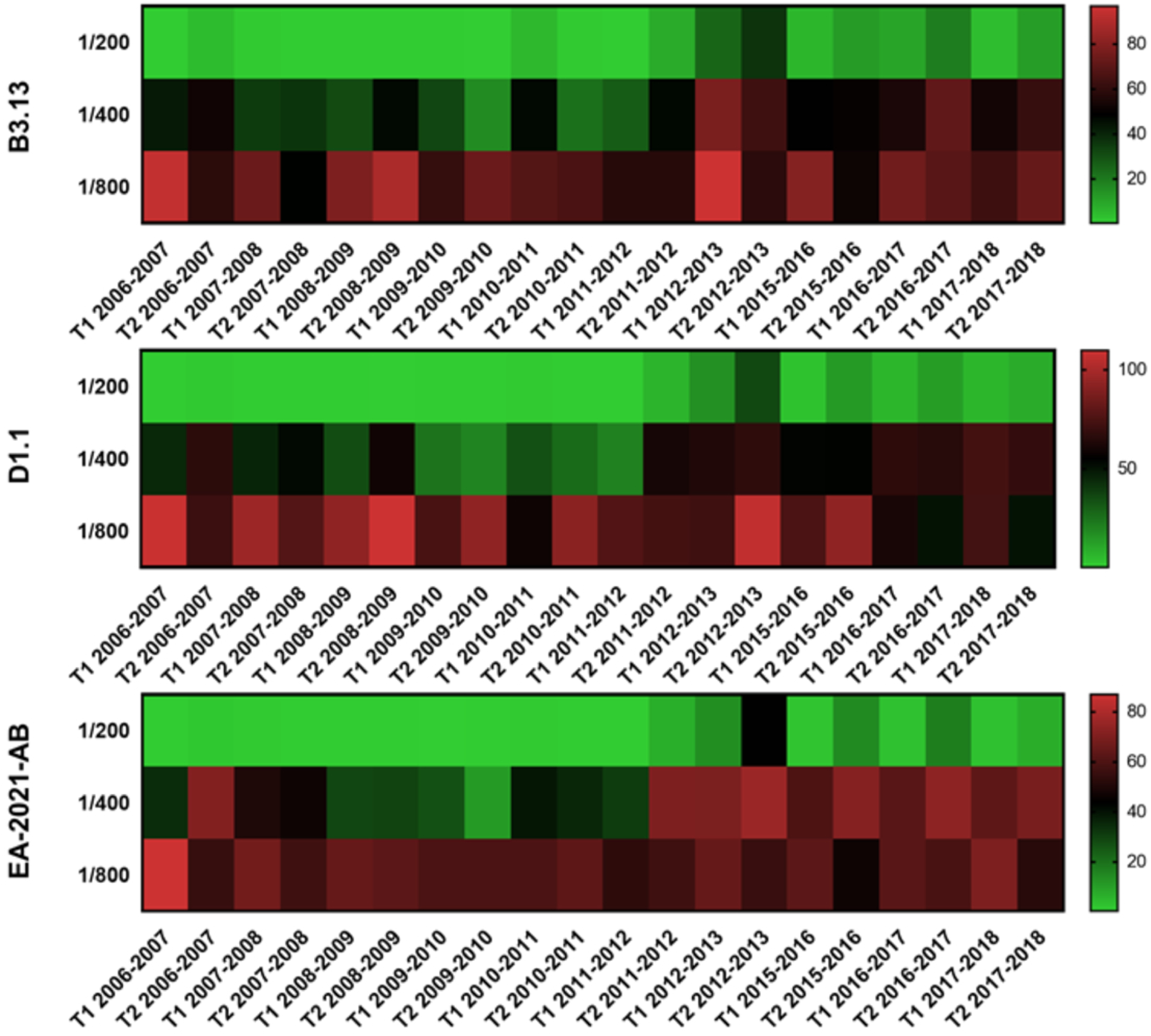
Detection of cross-neutralizing antibodies against H5N1 using NLuc reporter-virus assays. MDCK cells were seeded in 96-well plates (2×10^4^ cells/well, triplicates) and infected with 100-200 FFU of PR8(H5N1)-NLuc reporter viruses representing three clade 2.3.4.4b H5N1 genotypes: B3.13 (top), D1.1 (middle), and EA-2021-AB (bottom). After 1 h adsorption, the inoculum was replaced with fresh medium containing human serum samples at the indicated dilutions. Neutralization was quantified at 48 h post-infection by measuring NLuc activity in cell culture supernatants. Mock-infected wells served as background controls, and wells infected with the PR8(H5N1)-NLuc reporter viruses without serum were used to define the maximum (100%) infection/signal. Heatmaps display the NLuc readout (or derived neutralization values) for individual serum samples across seasons and timepoints (T1, pre-vaccination; T2, post-vaccination). Lower normalized NLuc signal indicates stronger neutralization (reduced viral replication), whereas higher NLuc signal indicates weaker neutralization.

## DISCUSSION

HPAI H5N1 viruses continue to pose a substantial threat to global public health and remain strong candidates for the next influenza pandemic. In this study, we evaluated basal heterosubtypic immunity against three genotypes (B3.13, D1.1, and EA-2021-AB) of clade 2.3.4.4b H5N1 IAVs and assessed the extent to which human seasonal IIV induce cross-reactive responses. This analysis was conducted using serum samples collected before and after IIV vaccination across ten consecutive influenza seasons, encompassing periods before and after the emergence of the 2009 pandemic H1N1 virus (pH1N1), and stratified across four adult age groups. To facilitate these analyses, we generated recombinant PR8-based viruses expressing the HA and NA of the three H5N1 genotypes, and NLuc, allowing sensitive evaluation of functional antibody responses using NLuc-based MN assays.

Traditionally, an HAI antibody titer of 1:40 has been considered indicative of approximately 50% protection against infection with seasonal H1N1 or H3N2 viruses (46). Although this threshold may not fully translate to zoonotic or potentially pandemic viruses such as H5N1, it is widely used in clinical trials of avian influenza vaccines and is supported by CBER/CHMP guidance. In the absence of alternative validated correlates, we therefore applied this benchmark to both HAI and MN assays. Using inactivated PR8(H5N1) viruses, our HAI data revealed very limited seroprotection and seroconversion against H5N1 across all seasons, with only a small fraction of individuals exhibiting HAI titers ≥1:40 and no significant increase following seasonal vaccination. These results contrast with the previous studies conducted with these serum samples, which showed a high seroprotection and seroconversion against both pH1N1 and H3N2 subtypes following seasonal influenza vaccination (57–59). These findings are consistent with the substantial antigenic divergence between the HA head domains and receptor-binding sites of avian H5N1 and human H1N1/pH1N1 viruses (**Figure 1**), which limits the induction of cross-reactive HA head–directed binding neutralizing antibodies. Moreover, the presence of different glycosylation patterns among pH1N1 and H5N1 viruses can be a reason for a lesser recognition of heterotypic antibodies (65, 66).

Given these limitations, we extended our analysis to NA-directed immunity. NA is the second major influenza surface glycoprotein and plays a critical role in viral egress and cell-to-cell spread (8, 24, 26). Antibodies that bind and inhibit NA activity can restrict viral infection, reduce viral shedding, and attenuate disease severity, even in the absence of strong HA-mediated neutralization. Importantly, both avian H5N1 and human H1N1/pH1N1 viruses encode N1 group 1 NAs, providing a mechanistic basis for cross-reactive anti-N1 antibody responses. Indeed, recent serological studies have demonstrated widespread functional anti-N1 activity against contemporary clade 2.3.4.4b H5N1 viruses in adult populations despite minimal or absent H5 HAI responses, with strong correlations observed between NAI titers against H5N1 N1 and pH1N1 N1 (67–74). Consistent with these reports, our data demonstrate that NA-directed cross-reactive antibodies are broadly present in human sera and can recognize the N1 proteins of multiple H5N1 genotypes, even when HAI titers remain low or undetectable. This pattern is biologically plausible because NA primarily exerts its function after viral entry by facilitating progeny virion release and promoting viral spread. As a result, NA-inhibiting antibodies may not prevent initial infection but can substantially limit viral egress and dissemination. In this context, NA immunity may be considered “quietly protective,” mitigating disease severity and potentially reducing transmission by restricting viral spread rather than blocking HA-mediated viral attachment and entry, which is the primary target of HAI assays.

Across influenza seasons, seasonal vaccination was associated with a modest increase in NA inhibition, with a more pronounced enhancement observed after 2009. A potential explanation for this temporal pattern is the widespread population exposure to the N1 NA introduced by pH1N1 through natural infection and in subsequent vaccine updates, which may have boosted or primed N1-specific immune memory (**Figure 1**). This priming could increase the likelihood of cross-reactive NA inhibition against heterologous HxN1 viruses, including HPAI H5N1. Supporting this hypothesis, pH1N1 infection has been shown to elicit broadly reactive human anti-N1 antibodies capable of inhibiting multiple HxN1 viruses and conferring protection in animal models (68, 70, 72–74). Notably, our NLuc-based MN assays, particularly the format that better capture post-entry inhibition, detected significant levels of neutralizing antibodies, further supporting a functional role for anti-NA antibodies present in human sera (**Figure 6**).

Importantly, the magnitude and quality of NA-directed responses induced by seasonal influenza vaccines may vary across seasons not only due to differences in population exposure histories, but also as a consequence of vaccine formulation, composition, and manufacturer. Influenza vaccine potency is standardized primarily on HA content, whereas NA content is not standardized and can vary substantially across manufacturers, production platforms, and vaccine lots, as well as in enzymatic activity and antigen stability. Consistent with this, differences in NA functional activity and in the induction of anti-NA responses have been reported between vaccine types, including split, subunit, and adjuvanted formulations. Therefore, the heterogeneity in NA inhibition observed across seasons in our study should be interpreted in the context of both immunological history and vaccine product characteristics. While ELLA provides a robust and reproducible functional readout of NA enzymatic inhibition *in vitro*, it is important to emphasize that NI titers represent a mechanistic surrogate rather than a direct correlate of *in vivo* protection. Although NA inhibition is frequently associated with reduced disease risk, it is not synonymous with clinical efficacy on its own. Thus, NA-directed immunity should be viewed as one component of a multifaceted protective response (24, 26, 28, 75).

Overall, our findings support the concept that seasonal influenza vaccination can induce cross-reactive immune responses that may confer limited but potentially meaningful protection at the population level against HAPI H5N1 viruses. These results are consistent with previous studies suggesting that seasonal IIV may provide partial benefit in the event of an H5N1 pandemic (16, 20, 21, 72, 73, 76, 77). However, our data also suggest that seasonal vaccination alone is unlikely to provide robust protection and may instead function primarily as an immunological priming step. In this scenario, subsequent boosting with H5N1-specific vaccines could elicit stronger, more specific, and long-lasting protective responses.

Seasonal IIV predominantly induce humoral immunity through B cell activation, and the precise role of memory B cells in mediating protection following natural exposure to AIV remains unclear. In addition, cellular immune responses, particularly T cell-mediated immunity, are likely to play a critical role in protection against novel IAV strains, including H5N1. Compared with IIV, live-attenuated influenza vaccines (LAIVs) have been shown to induce more robust cellular responses and to provide superior protection against heterologous or heterosubtypic viruses. Therefore, future studies evaluating the capacity of LAIVs to induce neutralizing and cross-reactive immune responses against HPAI H5N1 viruses with pandemic potential are warranted.

## Acknowledgments

We thank the Biodefense and Emerging Infectious Research Resources Repository (BEI Resources, for providing the following reagent: NR-59424). This project has been funded in part with Federal Funds from the National Institute of Allergy and Infectious Diseases, National Institutes of Health, Department of Health and Human Services, under contract no. 75N93021C00018 (NIAID Centers of Excellence for Influenza Research and Response, CEIRR). This project has been funded in part by the PID2023-146428OB-I00 funded by MCIN/AEI. This project has been funded in part by national funds from Gerencia Regional de Salud de la Junta de Castilla y León (GRS2557/A/22). This work was partially supported by a Texas Biomed Forum Award to A.M. Research on influenza to L.M.-S is supported by the American Lung Association (ALA).

## Contributions

Conceptualization: I.S.-M., J. M-E., L. M.-S., and A.N.

Methodology: I.S.-M., A.N., C.C.-G., A. M.-T., M. T.-A, A.M., J.S.-M., C.R.-C., I. B-M., and S. L.-B., L. M-S.

Formal Analysis: I I.S.-M., A.N., C.C.-G., A. M.-T., M. T.-A, A.M., J.S.-M., C.R.-C., I. B-M., and S. L.-B.

Funding Acquisition I.S.-M., and A.N.

Writing – Original Draft: I.S.-M., and A.N.

Writing-Review & Editing: I I.S.-M., A.N., C.C.-G., A. M.-T., M. T.-A, A.M., J.S.-M., C.R.-C., I. B-M., and S. L.-B., L. M-S.

All authors have approved the manuscript for publication.

